# Influence of delayed density and ultraviolet radiation on caterpillar baculovirus infection and mortality

**DOI:** 10.1101/2021.03.22.436482

**Authors:** Adam Pepi, Vincent Pan, Danielle Rutkowski, Vinay Mase, Richard Karban

**Affiliations:** Graduate Group in Ecology, University of California, Davis, CA 95616; Department of Entomology & Nematology, University of California, Davis, CA 95616

**Keywords:** Arctiinae, baculovirus, Bodega Marine Lab, disease dynamics, latitudinal gradient, population cycles

## Abstract

1. Infectious disease is an important potential driver of population cycles, but this must occur through delayed density-dependent infection and resulting fitness effects. Delayed density-dependent infection by baculoviruses can be caused by environmental persistence of viral occlusion bodies, which can be influenced by environmental factors. In particular, ultraviolet radiation is potentially important in reducing the environmental persistence of viruses by inactivating viral occlusion bodies.
2. Delayed density-dependent viral infection has rarely been observed empirically at the population level although theory predicts that it is necessary for these pathogens to drive population cycles. Similarly, field studies have not examined the potential effects of ultraviolet radiation on viral infection rates in natural animal populations. We tested if viral infection is delayed density-dependent with the potential to drive cyclic dynamics and if ultraviolet radiation influences viral infection levels.
3. We censused 18 Ranchman’s tiger moth (*Arctia virginalis*) populations across nearly 9° of latitude over two years and quantified the effects of direct and delayed density and ultraviolet radiation on baculovirus infection rates, infection severity, and survival to adulthood. Caterpillars were collected from each population in the field and reared in the laboratory. Baculovirus has not previously been described infecting *Arctia virginalis*, and we used genetic methods to confirm the identity of the virus.
4. We found that infection rate, infection severity, and survival to adulthood exhibited delayed density-dependence. Ultraviolet radiation in the previous summer decreased infection severity, and increased survival probability of the virus. Structural equation modelling indicated that the effect of lagged density on moth survival was mediated through infection rate and infection severity, and was 2.5 fold stronger than the effect of ultraviolet radiation on survival through infection severity. We successfully amplified polh, lef-8, and lef-9 viral genes from caterpillar samples, and BLAST search results confirmed that the virus was a nucleopolyhedrovirus.
5. Our findings provide clear evidence that delayed density dependence can arise through viral infection rate and severity in insects, which supports the role of viral disease as a potential mechanism, among others, that may drive insect population cycles. Furthermore, our findings support predictions that ultraviolet radiation can modify viral disease dynamics in insect populations, most likely through attenuating viral persistence in the environment.

## Introduction

Infectious disease plays an important role in population dynamics, potentially regulating populations and driving cycles. Many insect populations undergo dramatic cyclic fluctuations, and cyclic, delayed density-dependent disease outbreaks have been proposed as an explanation (Anderson & May, 1980, 1981). However, many other mechanisms have also been proposed to explain insect population cycles (Myers & Cory, 2013), particularly specialist parasitoids (Berryman, 1996). For some species of Lepidoptera, empirical evidence has suggested that baculoviruses are most likely the cause of cyclicity (Myers, 2000; Myers & Cory, 2013, 2016), though this has not been formally tested.

Delayed density-dependent feedbacks are required to drive cyclic population dynamics (Turchin, 2003). Detection of delayed density-dependence involving many ecological factors has been common (Turchin, 1990), and there have been several observations that viral epizootics follow high densities of some insect species (Cory & Myers, 2003; Fuxa, 2004; Myers, 2000; Myers & Cory, 2016). However, direct demonstrations of delayed density-dependent viral infection rate and infection-induced mortality are less common (Burthe et al., 2006; Fleming et al., 1986; Rothman, 1997). Observational studies over relatively small areas (>15km) using space-for-time substitutions of local densities have shown delayed density-dependent incidence of viral infection in voles (Burthe et al., 2006) and soil-dwelling hepialid caterpillars (Fleming et al., 1986). Experimental manipulation of western tent caterpillar (*Malacosoma californicum pluviale*) densities at the tree level showed delayed density-dependent infection rates by a nucleopolyhedrovirus (Rothman, 1997).

Besides host density, other aspects of the local environment may affect viral transmission and dynamics in the field (Cory & Myers, 2003). In particular, ultraviolet radiation has been shown to inactivate viruses of all kinds (Sagripanti & Lytle, 2007), including baculovirus occlusion bodies (Griego et al., 1985; Witt & Stairs, 1975). In field studies, examination of the presence of baculovirus occlusion bodies on shaded vs. unshaded foliage suggested that sunlight on leaves may inactivate viruses (Olofsson, 1988). A study of baculovirus transmission in forest tent caterpillars (*Malacosoma disstria*) found depressed transmission rates on forest edges as opposed to the forest interior, which was attributed to sunlight inactivating virus on leaves at forest edges (Roland & Kaupp, 1995). A field study of two strains of gypsy moth NPV showed variable resistance of virus to ultraviolet, in which the inactivation rate from ambient ultraviolet was greater in a more potent strain than a less potent one (Akhanaev et al., 2017). In another study of forest tent caterpillars using tree ring analyses, it was suggested that periods of weaker ultraviolet radiation increased the effect of forest tent caterpillar outbreaks on tree growth (Haynes et al., 2018). In this example, less ultraviolet radiation may have allowed better viral persistence and transmission and produced more severe caterpillar outbreaks.

In the present study, we examined the effects of host density, delayed-density dependence, and ultraviolet radiation on the survival, infection rate, and infection severity of a newly discovered baculovirus in Ranchman’s tiger moth (*Arctia virginalis*; Erebidae:Arctiinae). Ranchman’s tiger moth is a univoltine species with a long larval stage, during which it is a generalist herbivore (Karban et al., 2010). Caterpillars are parasitized by tachinid parasitoids, though caterpillars sometimes survive parasitism (English-Loeb et al., 1990). Parasitism has been found to have little role in population dynamics, leaving the observed oscillatory dynamics largely unexplained (Karban & de Valpine, 2010). Baculovirus is a dsDNA virus that persists outside of the host in the environment within a protective protein shell as an occlusion body (OB). Horizontal transmission can occur when caterpillars consume OBs on food sources, which dissolve in caterpillars’ alkaline guts and release infective virus particles (Vega & Kaya, 2012). Virus particles infect the caterpillar starting from the epithelial tissue and then move to other parts of the body, including the trachea, fat bodies, and hemolymph (Barrett et al., 1998), potentially causing death of the caterpillar.

To test the potential role of baculovirus in delayed-density dependent population dynamics in Ranchman’s tiger moth and the influence of ultraviolet radiation on the persistence and transmission of virus in moth populations, we censused 18 populations over two years along a gradient of latitude and ultraviolet radiation and reared caterpillars from these populations in the laboratory. To identify the mechanisms through which baculovirus affects moth population dynamics, we measured baculovirus infection rate, severity, and survival of caterpillars to adulthood. We analyzed the effect of caterpillar density and ultraviolet radiation at the study sites on viral infection and survival. We also tested if viral infection mediated the effects of caterpillar density and ultraviolet radiation on caterpillar survival using a structural equation model. In addition, we characterized the baculovirus identified in Ranchman’s tiger moths using sequence data and microscopy.

## Methods

### Description of system

In California, Ranchman’s tiger moths eclose as caterpillars in late summer, and feed and develop until the following spring. Pupation occurs in late spring, and adults emerge in early summer, mate and lay eggs during the summer. Early instars feed cryptically in litter and undergrowth until they become larger and climb higher onto vegetation in winter months. Caterpillars are generalists and feed on a variety of plants, with a preference for alkaloid-containing hosts (Karban et al., 2010), and occur mostly in open grassland, shrubland, or savannah. At the end of the larval stage, caterpillar movement increases, and caterpillars disperse away from patches to pupate (Grof-Tisza et al., 2014). Caterpillars also sometimes show symptoms of baculovirus infection, which include poor growth, inactivity, watery, opaque frass, regurgitation of milky fluid, and death. Caterpillars do not lyse at death; the integument of cadavers remains relatively intact. In dissections, viral occlusion bodies have been found in caterpillars, pupae, and adults. Therefore, it appears that viral infection may not always be lethal - in some cases, sublethal infections may persist in adults and pass on to the subsequent generation, although observations suggest that fecundity may be reduced in infected adults.

### Identification of baculovirus

To describe the virus, we removed infected fat tissue from four caterpillars from two sites, and amplified the *polh, lef-8*, and *lef-9* genes using oligonucleotides with universal primers attached for the *lef-8* and *lef-9* genes using PCR (following Jehle et al., 2006). PCR products were then sequenced. We also smeared fat body tissue and stained with buffalo black and examined under a light microscope with a 400x oil immersion lens. Sequencing results were poor for polh due to short DNA fragments, but we conducted a BLAST search on NCBI using lef-8 and lef-9 gene sequences.

### Censuses

Caterpillars were counted visually along 4 m wide linear transects of varying length (78-300m) at 18 sites along a ~1000 km length adjacent to the Pacific coast in California, Oregon, and Washington (Figure 1). Transects were surveyed in 2019 and 2020, starting from 1 March through 30 May. We visited sites in each year from south to north, in accordance with the phenology of caterpillar development so that caterpillars were surveyed when most were 4^th^ or 5^th^ instars (sampling dates in Table S1). Transects were surveyed at a constant walking speed and all by the same observer (A. Pepi). Density was estimated from caterpillar counts over the transect area (4m x length). During the second year, sites were revisited within ten days of the calendar date of the first visit; however, this species is a slow-growing caterpillar (roughly ten month development period), so density estimates were likely not overly sensitive to small deviations in sampling date. In 2020, up to ca. 30 caterpillars per site (or less if fewer were found) were collected haphazardly and brought back to the laboratory for rearing.

**Figure 1.**
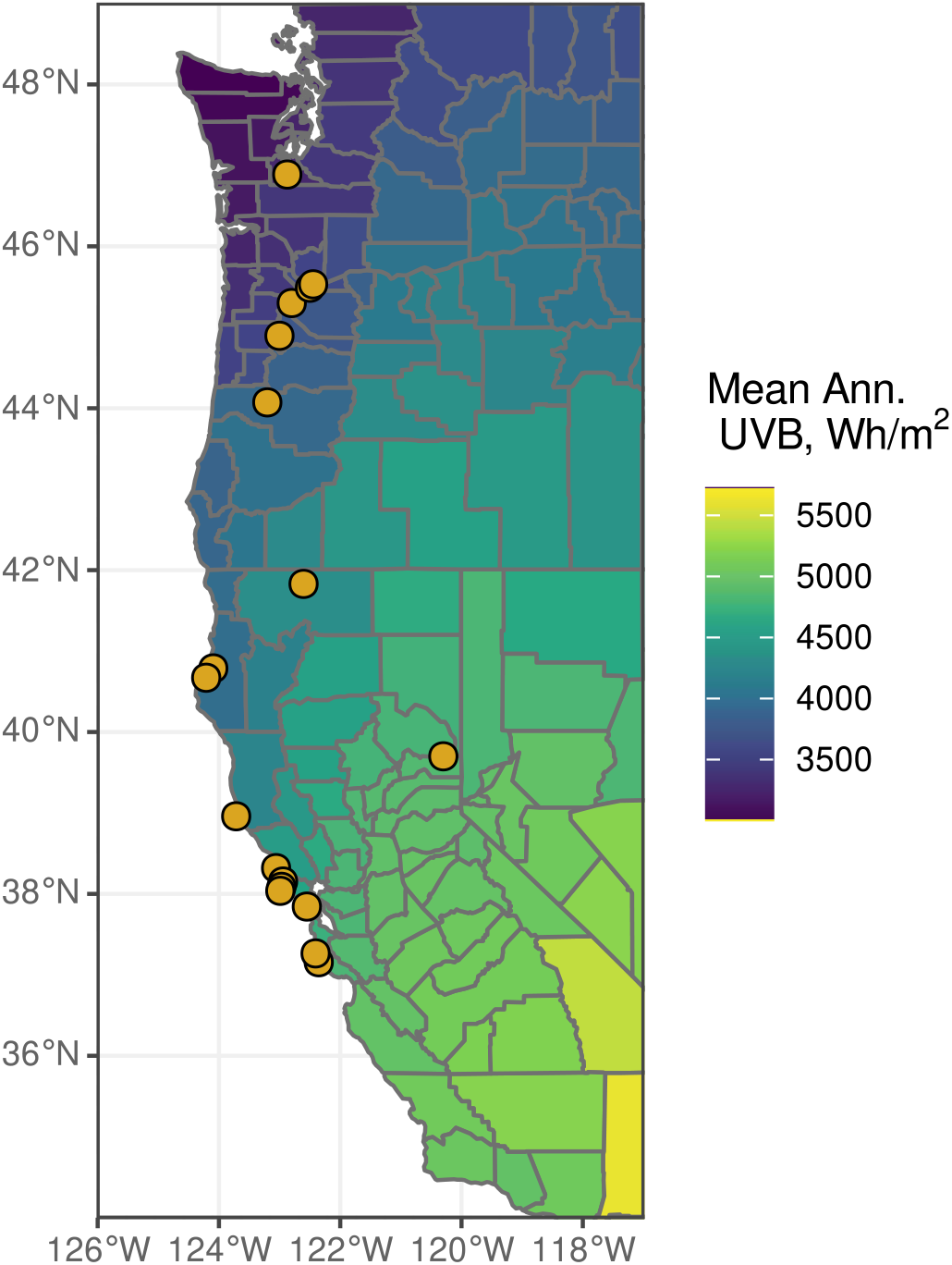
Map of study site locations in Washington, Oregon, and California (orange points), overlaid on a map of mean annual UV-B radiation aggregated at the county level, in Wh/m^2^ (Tatalovich et al., 2006).

### Rearing

Caterpillars were reared individually in 6 oz plastic souffle cups with fabric lids and kept in an incubator at 18 °C and 75% RH. Caterpillars were fed every 3-4 days with washed organic romaine lettuce and yellow bush lupine (*Lupinus arboreus*) leaves collected from a part of Bodega Marine Reserve without *Arctia virginalis*. Parasitoids that emerged from caterpillars were counted and identified to family. Caterpillars that died during rearing were frozen at −20 °C for subsequent dissection, except for those that clearly died due to emerged parasitoids. Caterpillars were reared until pupation (~50 days), and pupae were placed together by site of origin (no more than 10 per site) in 30 cm x 30 cm flight cages and allowed to emerge as moths, at room temperature. Pupae and moths were reared until one month after the last moth emerged and sprayed once or twice weekly with water to prevent desiccation. Adults do not feed. Dead pupae and moths were stored dry at room temperature for subsequent dissection. Individuals that reached adulthood fully formed were counted as having survived; individuals that died as caterpillars or failed to emerge from pupae or expand wings were counted as not surviving.

### Dissection and viral assays

After death, each individual was dissected to assess infection status. Two to six tissue samples per insect were taken from the abdomen of adults and pupae or fat bodies of caterpillars, broken up with forceps, smeared on a glass slide with water, and examined under a light microscope at 200x magnification with phase contrast. Visible occlusion bodies (OB) (<0.15μm) were confirmed by adding a drop of 1 M NaOH to the slide, which dissolves the viral protein coat, turning the OB transparent (Lacey & Solter, 2012)(Figure S1). Tissue samples were taken until a positive NaOH test was obtained; if no virus was found after six tissue samples, individuals were classified as uninfected.

Infection severity was rated as uninfected, low, medium, high, or very high, based on the number of samples required to discover virus, the density of OBs in tissue smear samples, and the coloration of hemolymph and fat body (Figure S2). Low severity infections took many samples (4-6) to identify and had low densities of OBs. Medium severity infections took fewer samples (1-2) to identify and had moderate densities of OBs. High severity samples had infections in all samples with high densities of OBs, with organ systems in the body still intact and identifiable. Very high severity infections had infections in all samples with very high densities of OBs, with internal organ systems unidentifiable.

### Ultraviolet irradiation

We calculated the cumulative ultraviolet radiation in 2019 for each site that OBs are likely to have experienced. Daily values for UV radiation, specifically ‘Erythemal Daily Dose’ in Watts/m^2^ from OMI/Aura satellite data were accessed from NASA GES DISC (Hovila et al., 2014). Values were extracted for each sampling location and the average value calculated from 360 to 180 days before the collection date of caterpillars (~March-September at most sites; see Table S1 for collection dates). This timing coincided with the season during which viral OBs would have persisted in the environment outside of hosts.

### Statistical analysis

First, we analyzed the density-dependence of infection rate (proportion of infected individuals) and the influence of ultraviolet radiation on infection rate. The proportion of all individuals (N=208) that were infected was analyzed in response to log caterpillar density per m^2^ from transects in 2019 (density in the previous year) and 2020 (density in the same year) using beta-binomial generalized linear mixed models with site as a random effect. To test for the effect of ultraviolet radiation, we implemented a model with both ultraviolet radiation and log density, to control for the effect of density on infection rate.

Second, we analyzed the density-dependence of infection severity, and the influence of UV radiation on infection severity. We analyzed infection severity (N=198 with severity rated) in response to log density (same year and previous year) using cumulative link mixed models with site as a random effect. As before, to test for the effect of UV radiation, we implemented a model with both UV radiation and log density, to control for the effect of density on infection severity. To test for effects of density and UV on infection severity independent of infection status, we implemented models using only infected individuals (N=184). This allowed us to test whether different processes influenced infection rate vs. infection severity. We implemented cumulative link mixed models with only log density, and both log density and UV radiation as predictors as before.

Third, we tested for density-dependent survival and effects of UV radiation on survival. We analyzed survival to emergence as fully formed moths (N=208) in response to caterpillar density, tachinid parasitoid load, and UV radiation in a beta-binomial generalized linear mixed model with site as a random effect.

Lastly, we tested our structural hypothesis that viral infection and infection severity were the mechanisms through which lagged density and UV radiation affected survival in moth populations. There are multiple possible pathways through which density, UV radiation, infection rate, and infection severity might influence survival and thus population dynamics. To test our specific structural hypothesis, we constructed a piecewise structural equation model which tested whether infection rate and infection severity mediated the effects of UV radiation and caterpillar density on survival. To do this, we transformed infection severity into a continuous variable between 0 and 1, by converting to numerical categories from 1-5, dividing by 5, and subtracting 0.1. We constructed 3 component submodels of our structural equation model: a beta-binomial model of infection rate predicted by log density and UV radiation, a beta model of infection severity rate predicted by infection rate, log density and UV radiation, and a beta-binomial model of survival predicted by infection rate and infection severity (Figure 5). All sub-models included site as a random effect, and only individuals that were not parasitized and were scored for infection severity were included in the analysis (N=198). Standardized coefficients were calculated using the latent-theoretic method (Grace et al., 2018).

Beta-binomial and beta generalized linear mixed models were fitted using glmmTMB (Brooks et al., 2017). Cumulative link mixed models were fitted using the ordinal package (Christensen, 2019). Piecewise structural equation models were fit using piecewiseSEM v. 1.2.1 (Lefcheck, 2016), and component models were fit using glmmTMB. UV radiation was logged and scaled to improve model convergence. We checked for multicollinearity in models using VIFs; multicollinearity was low to moderate in all models (<6) except for same year density in models of infection severity (VIF from 10.5-11.5). Since we included same year density as a control variable, we kept these models as is, despite high VIFs. Plots were generated using ggplot2 (Wickham, 2009), ggeffects (Lüdecke, 2018), and viridis (Garnier, 2018). All analysis and plotting were conducted in R v. 3.6.3 (R Development Core Team 2020).

## Results

### Description of virus

We successfully amplified *polh, lef-8* and *lef-9* genes with PCR from four infected individuals. The BLAST search resulted in top matches of *Perigonia lusca* NPV and *Anticarsia gemmatalis* NPV using lef-8 and lef-9 genes respectively (Table S2, S3). This result, along with size of occlusion bodies under the microscope, indicated that the *Arctia virginalis* baculovirus was a nucleopolyhedrovirus since both BLAST matches were alphabaculoviruses (NPV).

### Infection rate

The probability of infection increased with log density in the previous year in models of infection rate (Table 1, Figure 2a). Infection rate had little relation to log density in the current year or to UV radiation over the season when environmental OBs could have been exposed (Table 1, Figure 2b).

**Figure 2.**
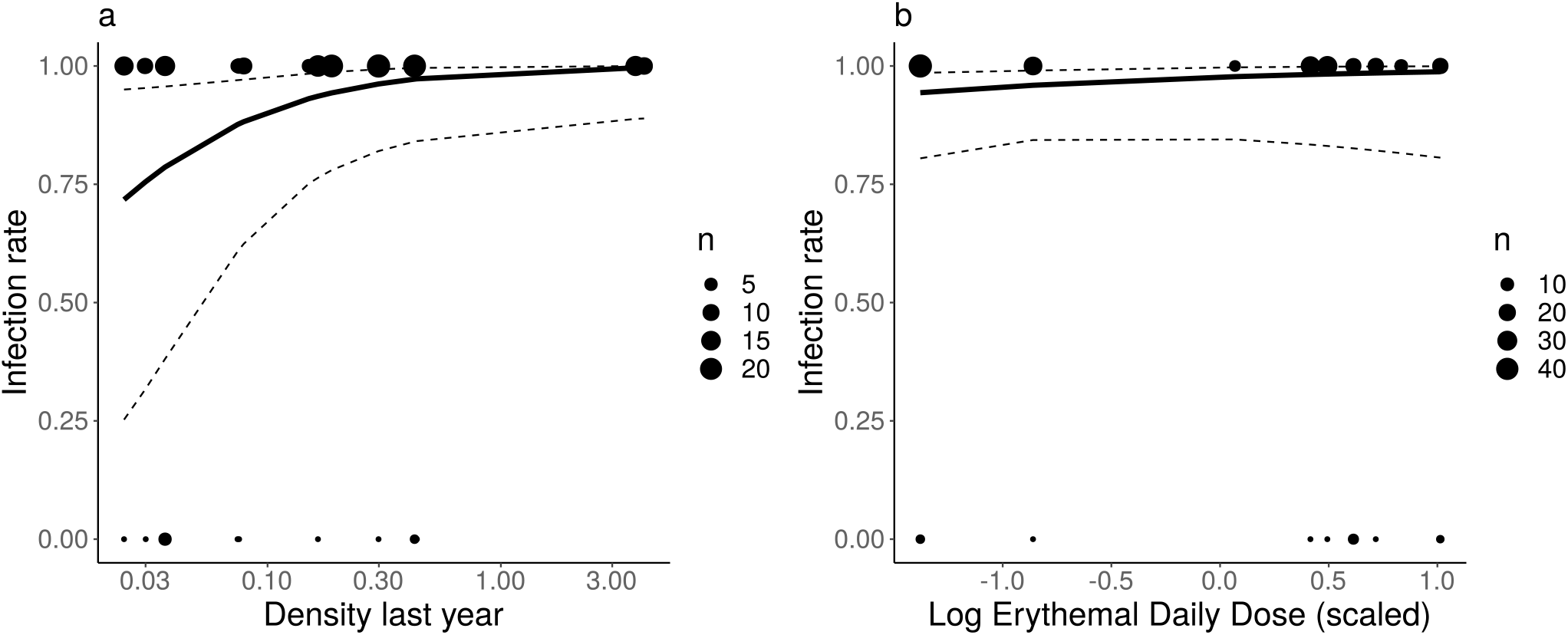
Infection rate in response to (a) density in the previous year and (b) ultraviolet radiation as scaled log erythemal daily dose in the summer before collection (2019) from beta-binomial generalized linear mixed models.

**Table 1.**
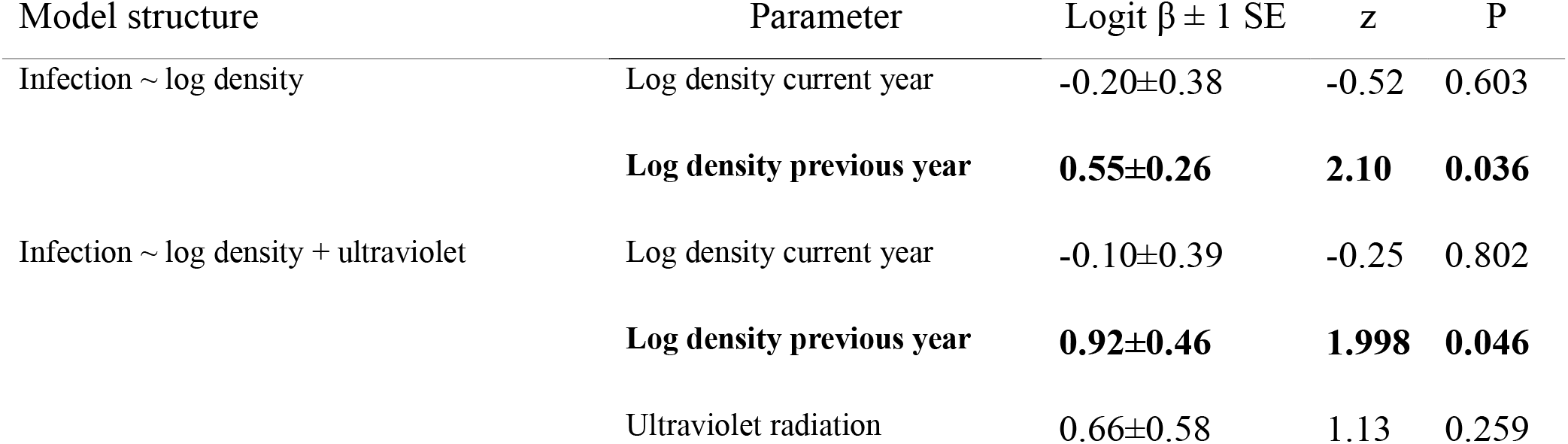
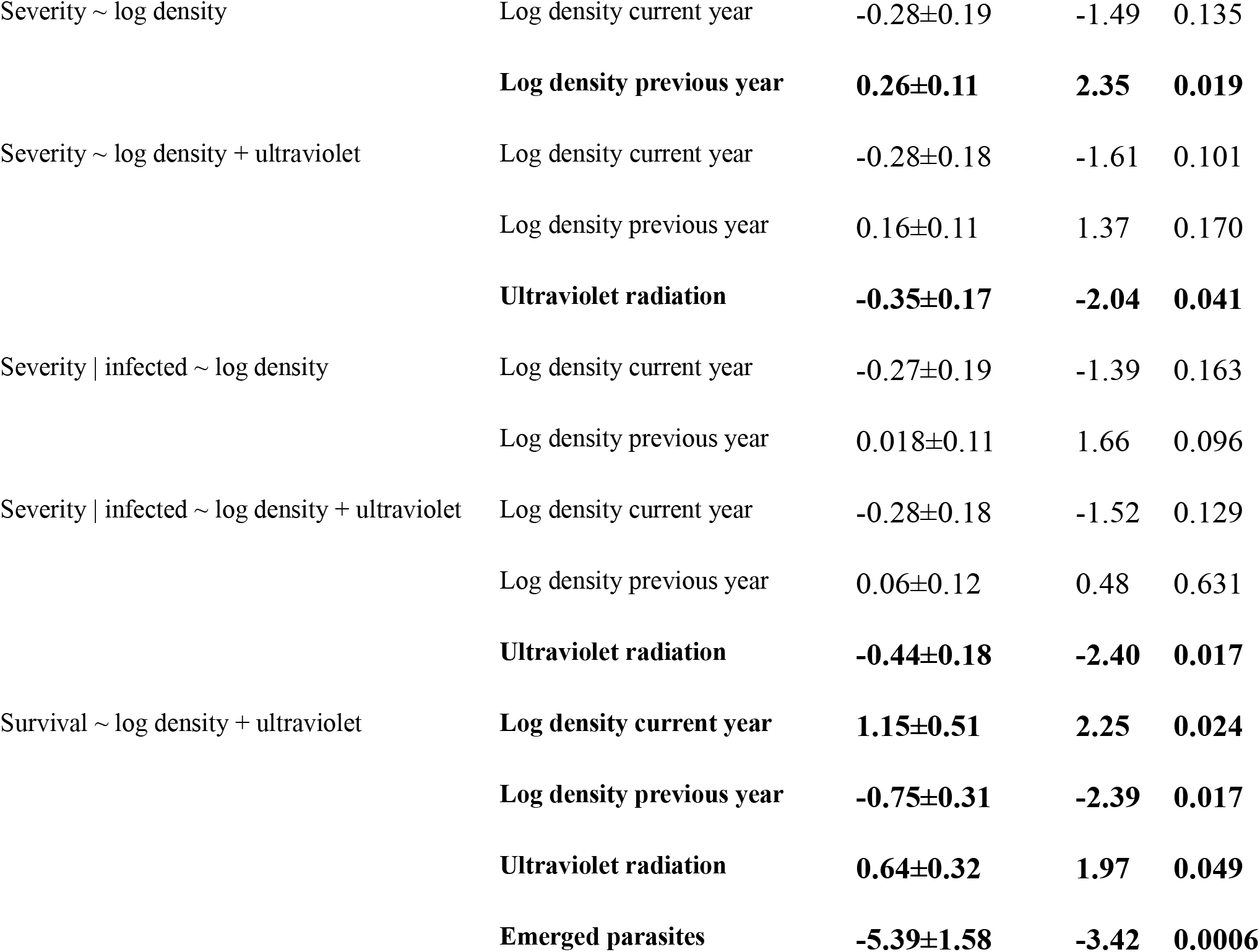
Results of univariate models.

### Infection severity

Log density in the previous year increased infection severity in models of infection severity that included both infected and uninfected caterpillars but this effect was weakened when UV radiation was included in the model (Table 1, Figure 3a,b). This effect of UV radiation may have been due to low power or to a negative correlation between log density in the previous year and UV radiation (Pearson’s r: −0.53). Ultraviolet radiation decreased infection severity when included (Table 1, Figure 3c,d). In models that included only infected caterpillars, density in the current and previous years had little effect on infection severity, and UV radiation had a negative effect on infection severity (Table 1).

**Figure 3.**
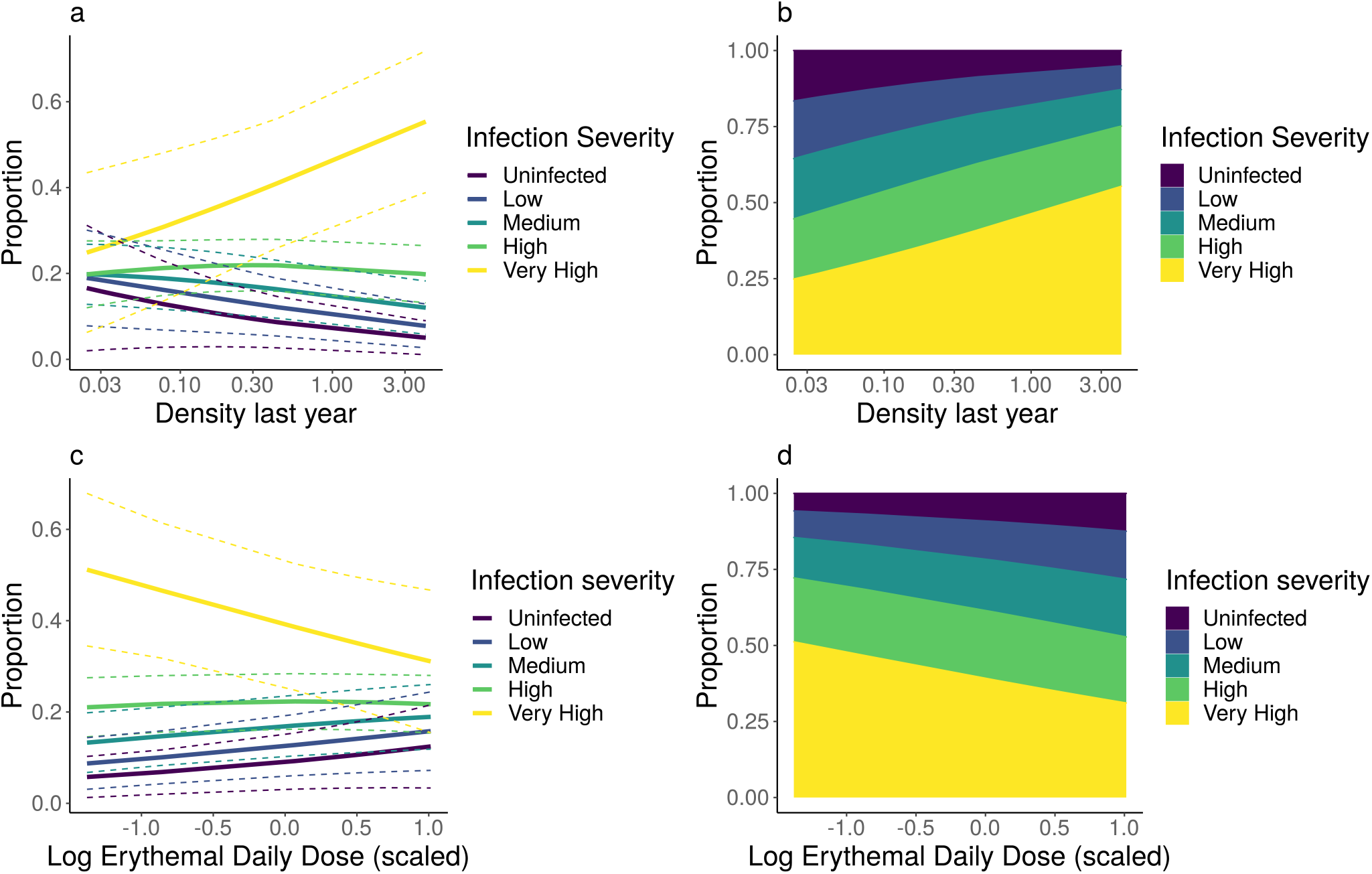
Infection severity in response to (a-b) density in the previous year, and (c-d) ultraviolet radiation as scaled log erythemal daily dose in the summer before collection cumulative link mixed models. The left column (a,c) shows model-predicted rates of infection at each severity class with 95% confidence intervals, and the right column (b,d) shows the proportion of the total population predicted to be infected at each severity class by color.

### Survival to adult emergence

Survival to adult emergence decreased with log density in the previous year and with the number of emerged parasites. Survival increased in response to density in the current year, and in response to UV radiation (Table 1, Figure 4).

**Figure 4.**
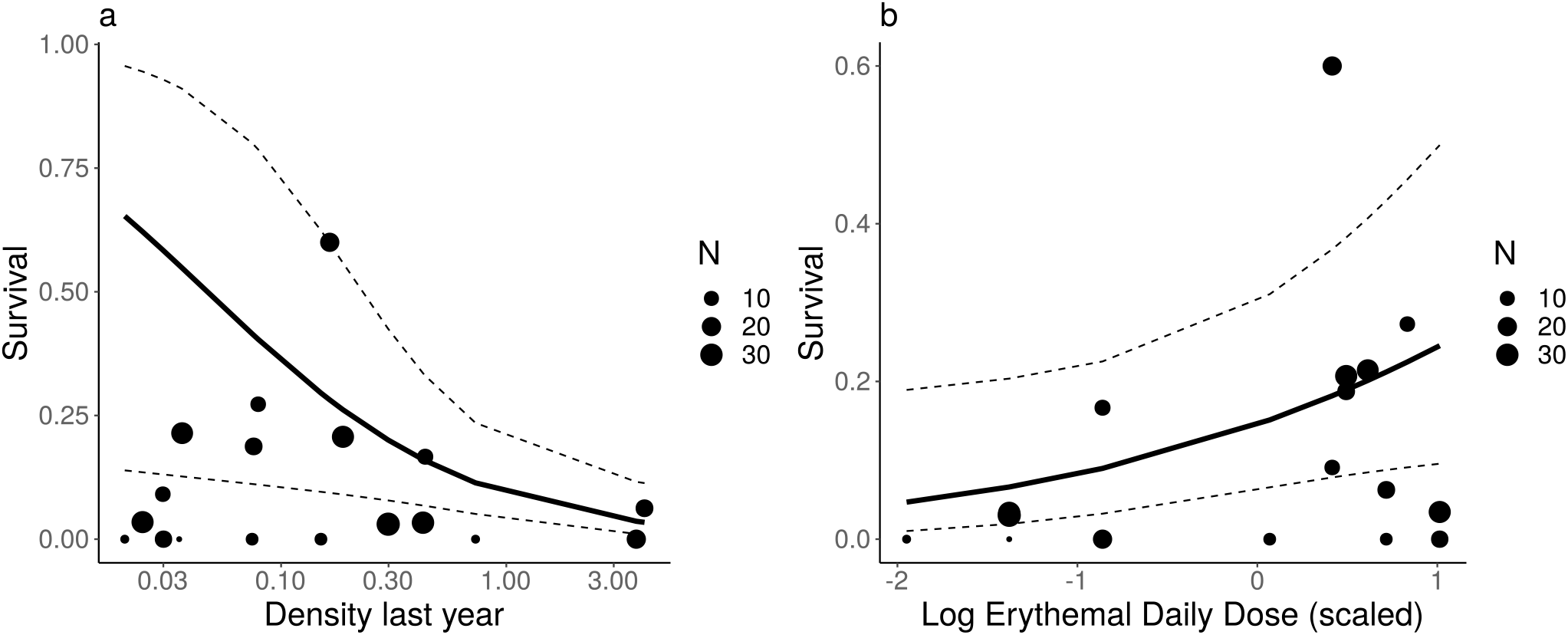
Survival to adult emergence in response to (a) density in the previous year, and (b) ultraviolet radiation as scaled log erythemal daily dose in betabinomal generalized linear mixed models. The size of each point is scaled to the number of individuals reared from each site.

### Structural equation model

We hypothesized a causal model linking UV radiation and disease to caterpillar survival (Fig. 5). Shipley’s d-separation test (Shipley, 2000) indicated that our structural equation model was correctly specified (Fig. 5, Fisher’s C = 7.38, df = 6, P = 0.288), meaning that there were no missing paths in our causal model. The model results showed that the negative effect of log density in the previous year on survival was mediated by infection rate and infection severity and that the positive effect of UV radiation on caterpillar survival was mediated by infection severity (Fig 5). The relative indirect negative effect size of log density in the previous year was ~2.5x greater than the positive effect of UV radiation on survival (−0.085 vs. 0.035, calculated by multiplying path coefficients).

**Figure 5.**
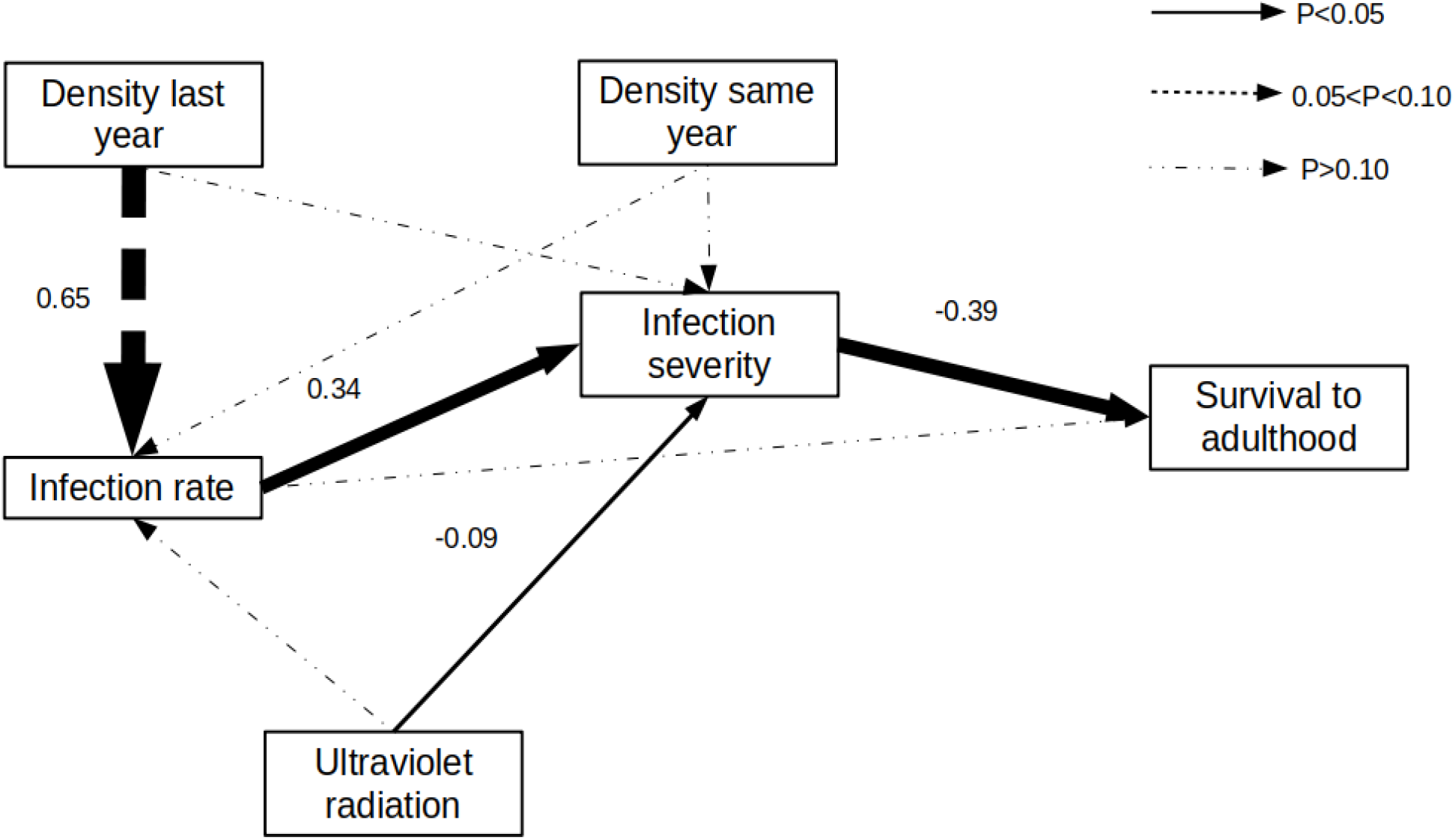
Path diagram of piecewise structural equation model. The size of the arrows is proportional to the standardized path coefficient (shown next to lines), and line type indicates P-value of the path parameter.

## Discussion

Overall, our results show negative delayed density-dependent effects of the virus on caterpillar population dynamics. Log density during the previous year increased infection rate and infection severity and decreased survival to adult emergence. The effect of delayed density on both infection rate and infection severity in univariate models and the result of the structural equation model suggest that the effect of density on survival acts through infection severity as well as infection rate. This is likely due to dose-dependent infection severity, in which caterpillars that are exposed to more viral occlusion bodies develop more severe infections that result in death (Cabodevilla et al., 2011; Eberle et al., 2008; Matthews et al., 2002).

In contrast, log density in the current year, representing direct density-dependence, had no effect on infection rate, a marginal negative effect on infection severity, and a positive effect on survival to adult emergence. Though inconsistent, this positive effect may have been due to ascertainment bias: sites that had larger viral outbreaks may have already declined in density over the course of the larval season (summer 2019-spring 2020). Therefore, density at the time of collection had potentially a reversed relationship to environmental OB density: sites that had declined from earlier high densities would have higher levels of OB, resulting in more severe infections and a higher probability of death. Overall, our results indicated negative delayed density dependence and weak or positive direct density-dependence consistent with the second-order, oscillatory population dynamics observed in the system over the long term (Pepi et al., 2021).

Ultraviolet radiation by contrast had no detectable effect on infection rate but reduced infection severity and increased caterpillar survival. The reduction in infection severity was even greater when only infected individuals were considered. The effect of ultraviolet radiation on infection severity and survival but not infection rates may represent dose-dependent effects of exposure to OBs. Specifically, at sites with higher ultraviolet radiation, more viral OBs were likely inactivated before caterpillars could be exposed to them, so caterpillars received smaller doses of virus and thus became less severely infected and were more likely to survive.

The observational nature of our study gives it some strengths and weaknesses. Our study is one of few to show delayed-density dependent viral infection in the field, and the only study to show effects of radiation on viral infection in a natural population of insects. Our results provide validation in a field system to laboratory studies of virus dynamics and inactivation effects of UV radiation on viruses (Bjørnstad et al., 1998; Witt & Stairs, 1975). Despite highly variable conditions between sites, which were spread across a gradient of over 1000 km, we were able to detect clear effects of delayed density on infection rate, severity, and survival to adulthood, and of ultraviolet radiation on infection severity and survival to adulthood. We were also able to use structural equation modelling to show that infection rate and severity mediate the effect of delayed density and UV radiation on survival and strengthen the inferences made from our observational study.

Density in the previous year was a strong predictor of infection rates in populations (Fig 2). This finding makes sense because there must have been a large enough population of hosts in the previous year for the disease to spread and produce sufficient inoculum to persist into the current year. Density in the previous year also affected infection severity (Fig 3), but this effect became much weaker when UV radiation was included in models and even weaker when only infected caterpillars were considered. In the structural equation model, infection severity was well explained by infection rate because individuals must be infected to have high infection severity. This result may be due to the conflation of infection rate and infection severity in the model since the same dose-dependent mechanism may have caused them. However, in all models of infection severity that included both UV radiation and delayed density, the effect of delayed density, beyond whether individuals were infected or not, was weaker than the effects of UV radiation. Thus, when effects on infection severity were detected, they may have in fact been generated by the spurious negative relationship between delayed density and UV radiation. In contrast to infection severity, UV radiation had no significant effect on infection rate. Ultimately, both delayed density and UV radiation had strong and opposing effects on survival to adulthood, though the effect of density in the previous year was much greater in magnitude. The effects of delayed density on survival were mediated through infection with limited evidence of direct effect on infection severity. In contrast the effects of UV radiation on survival were mediated entirely through effects on infection severity.

A possible explanation for the finding that infection rates were primarily determined by host density in the previous year but not by ultraviolet radiation is that viral infection may be determined in part through vertical transmission (Burden et al., 2002; Cabodevilla et al., 2011; Cory, 2015; Cory & Myers, 2003). In particular, vertical transmission is likely to occur in *A. virginalis* through persistent sub-lethal infections. We regularly observed OB in egg samples in adult dissections, and thus eggs of adults with sublethal infections are likely contaminated with virus. Vertical transmission represents a mechanism through which density in the previous year might influence infection rates in the following year but is not subject to influence from ultraviolet radiation. In this way, vertical transmission may maintain higher infection rates after high-density years with epizootics, even if UV radiation attenuates inoculum in the environment (Cabodevilla et al., 2011). Sublethal infections may represent another means by which baculovirus affects population dynamics by imposing fitness costs on adults (Cabodevilla et al., 2011; Matthews et al., 2002; Rothman, 1997), although we did not measure this in the present study.

In the present study, we also give the first identification of a baculovirus pathogen of *Arctia virginalis* (AvNPV). Based on lef-8 and lef-9 genes, we conducted a BLAST search which identified this virus as a nucleopolyhedrovirus, which is consistent with the appearance of stained occlusion bodies under a light microscope (personal communication, J.S. Cory). However, the likely placement of AvNPV on the baculovirus phylogeny was not clear, since the two matched species, *Perigonia lusca* NPV and *Anticarsia gemmatalis* NPV were in the NPV clades II and I, respectively (Thézé et al., 2018). Further investigation and phylogenetic work will be necessary to identify the phylogeny of *Arctia virginalis* NPV. Caterpillars infected with AvNPV exhibited unusual symptoms, such as failure to lyse, and high survival rates despite heavy infection, which is atypical for NPV infections. In most well studied species, NPV infection leads to death relatively rapidly, and causes caterpillars to lyse. In the present case, this could be due to differences between AvNPV and other NPVs, such as in enzymes that decompose the insect integument, or host differences. *Arctia virginalis* caterpillars are longer lived and more physically robust than other better-studied lepidopteran baculovirus hosts, such as *Lymantria dispar* or *Spodoptera* spp.. Further studies should investigate some of the mechanisms of host-pathogen interactions between AvNPV and *Arctia virginalis* that lead to these unique characteristics, and their consequences for viral transmission and host-pathogen dynamics.

In summary, we demonstrated population-level delayed density-dependent effects on viral infection rate, infection severity, and survival to the reproductive stage, showing how viral infection may drive cyclic dynamics in insects. We also showed for the first time that UV radiation may influence disease dynamics and ultimately population dynamics of insects, through decreased infection severity and increased insect survival rates. This provides support for the proposal that viral epizootics may be an important mechanism driving cyclic dynamics of insects and Lepidoptera in particular (Myers & Cory, 2013). Our findings also suggest that UV radiation may be an important factor to consider as a driver of insect viral disease in the context of global change. Long-term changes in atmospheric transmittance of solar radiation, or “global dimming and brightening,” have been observed and are potentially anthropogenic in origin (Wild, 2016). Such long-term changes may have the potential to influence insect viral disease and population dynamics through changing levels of UV attenuation of viral inoculum (Haynes et al., 2018).

## Acknowledgements

We would like to thank Marcel Holyoak, Jenny Cory, and Jay Rosenheim for helpful comments on the manuscript. We would also like to thank California State Parks, Ben Becker at Point Reyes National Seashore, Kristen Ward at Golden Gate National Recreation Area, Jackie Sones at Bodega Marine Reserve, Brendan Leigh at Humboldt Bay NWR, Karlee Jewell at North Coast Landtrust, Diane Steeck at the City of Eugene, Grey Wolf at the City of Salem, Oregon Metro Parks, and Christian Haaning at the City of Portland for assistance and access to field sites. We would also like to thank Harry Kaya and Jenny Cory for assistance with identifying the baculovirus, Jay Rosenheim for the use of incubators, Neal Williams for the use of compound microscopes, and Claire Beck and Jasmine Daragahi for help with rearing caterpillars. The research was funded by NSF-LTREB-1456225 and an NSF REU supplement (DEB-2018169).

## Author contributions

AP, VP, and RK conceived the study. AP and VP collected the data and conducted the analyses. DR and VM conducted genetic work. AP wrote the manuscript. All authors contributed critically to the drafts and approved final publication.

## Data Availability

Data will be archived on KNB upon acceptance.

## Supplement

**Table S1.**

| Site | Latitude | Longitude | Number reared | Date Collected |
| --- | --- | --- | --- | --- |
| Franklin Point, CA | 37.155 | -122.35 | 29 | 3/1/2020 |
| Pescadero Marsh, CA | 37.265 | -122.41 | 16 | 3/1/2020 |
| Golden Gate NRA, CA | 37.844 | -122.55 | 28 | 3/13/2020 |
| Point Reyes South, CA | 38.038 | -122.99 | 15 | 3/13/2020 |
| Point Reyes North, CA | 38.088 | -122.97 | 29 | 3/13/2020 |
| Point Reyes Kehoe, CA | 38.153 | -122.94 | 2 | 3/13/2020 |
| Bodega Marine Reserve, CA | 38.319 | -123.06 | 16 | 3/27/2020 |
| Manchester State Park, CA | 38.959 | -123.72 | 6 | 3/27/2020 |
| Loyalton, CA | 39.700 | -120.30 | 6 | 5/30/2020 |
| Humboldt NWR, CA | 40.670 | -124.21 | 20 | 4/10/2020 |
| Freshwater Farms, CA | 40.786 | -124.09 | 11 | 4/10/2020 |
| Klamath River, Yreka, CA | 41.829 | -122.61 | 11 | 4/10/2020 |
| Checkermallow Access, OR | 44.072 | -123.20 | 12 | 4/11/2020 |
| Fairview Wetland, OR | 44.895 | -123.00 | 20 | 4/11/2020 |
| Graham Oaks, OR | 45.299 | -122.80 | 30 | 4/11/2020 |
| Powell Butte, OR | 45.488 | -122.50 | 33 | 4/11/2020 |
| Salish Ponds, OR | 45.531 | -122.45 | 1 | 4/11/2020 |
| West Rocky Prairie, WA | 46.887 | -122.87 | 2 | 4/12/2020 |

**Table S2.** Blast results for viral genes amplified from infected caterpillars.

| Sample ID | Gene | Matched taxon | Max Score | Total Score | Query cover- age | E value | Percent identity |
| --- | --- | --- | --- | --- | --- | --- | --- |
| Loyalton 6 | lef-8 | Perigonia lusca single NPV | 80.5 | 80.5 | 9% | 2E-10 | 87.69% |
| Loyalton 6 | lef-9 | Anticarsia gemmatalis MNPV | 130 | 130 | 84% | 9E-26 | 77.16% |
| Loyalton 24 | lef-8 | Perigonia lusca single NPV | 78.7 | 78.7 | 9% | 9E-10 | 87.69% |
| Loyalton 24 | lef-9 | Anticarsia gemmatalis MNPV | 130 | 130 | 84% | 9E-26 | 77.16% |
| Graham Oaks 10 | lef-8 | Perigonia lusca single NPV | 78.7 | 78.7 | 9% | 9E-10 | 87.69% |
| Graham Oaks 10 | lef-9 | Anticarsia gemmatalis MNPV | 124 | 124 | 84% | 4E-24 | 76.86% |
| Graham Oaks 20 | lef-8 | Perigonia lusca single NPV | 78.7 | 78.7 | 9% | 9E-10 | 87.69% |
| Graham Oaks 20 | lef-9 | Anticarsia gemmatalis MNPV | 117 | 117 | 84% | 7E-22 | 76.42% |

**Table S3.**
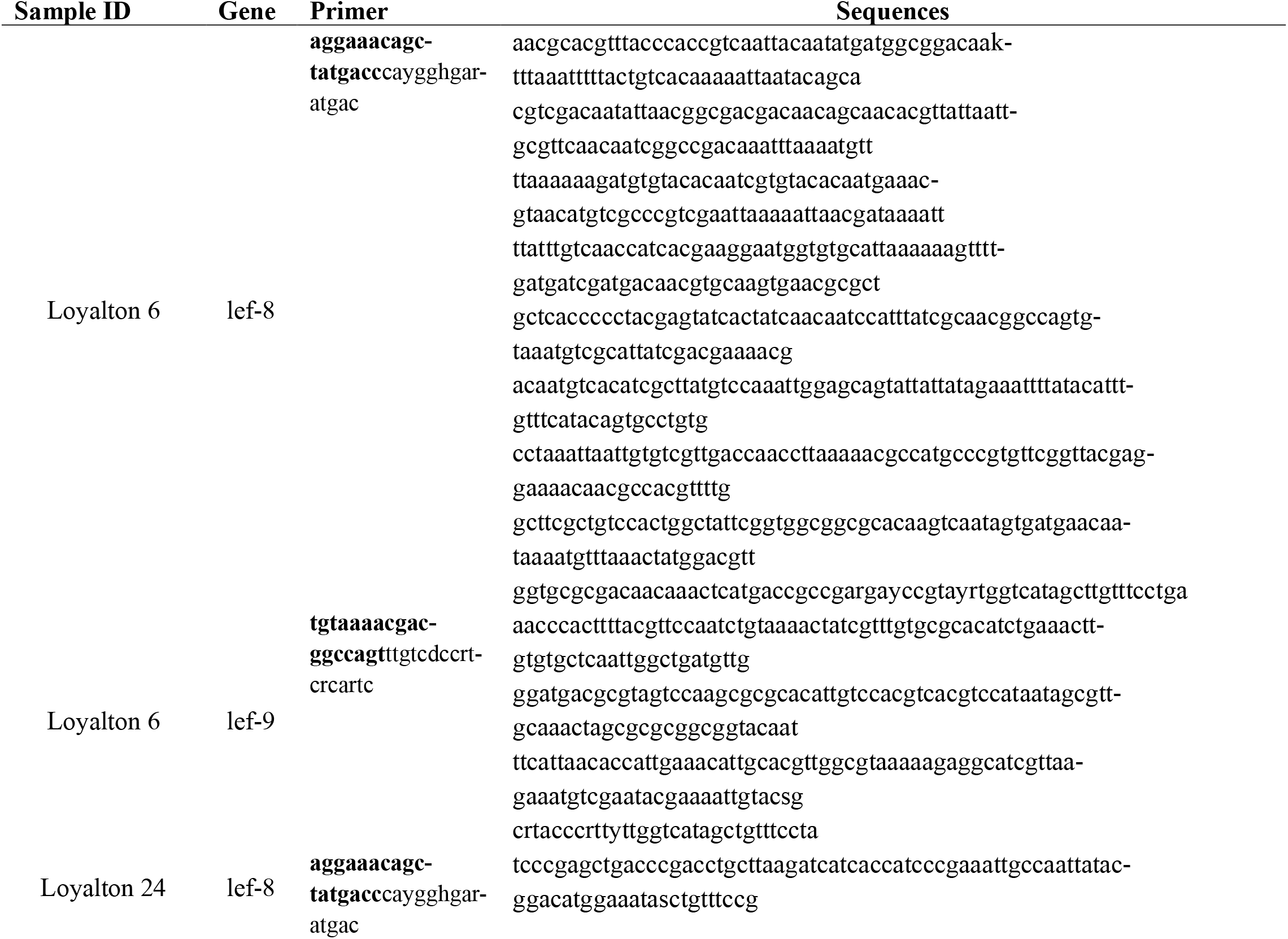
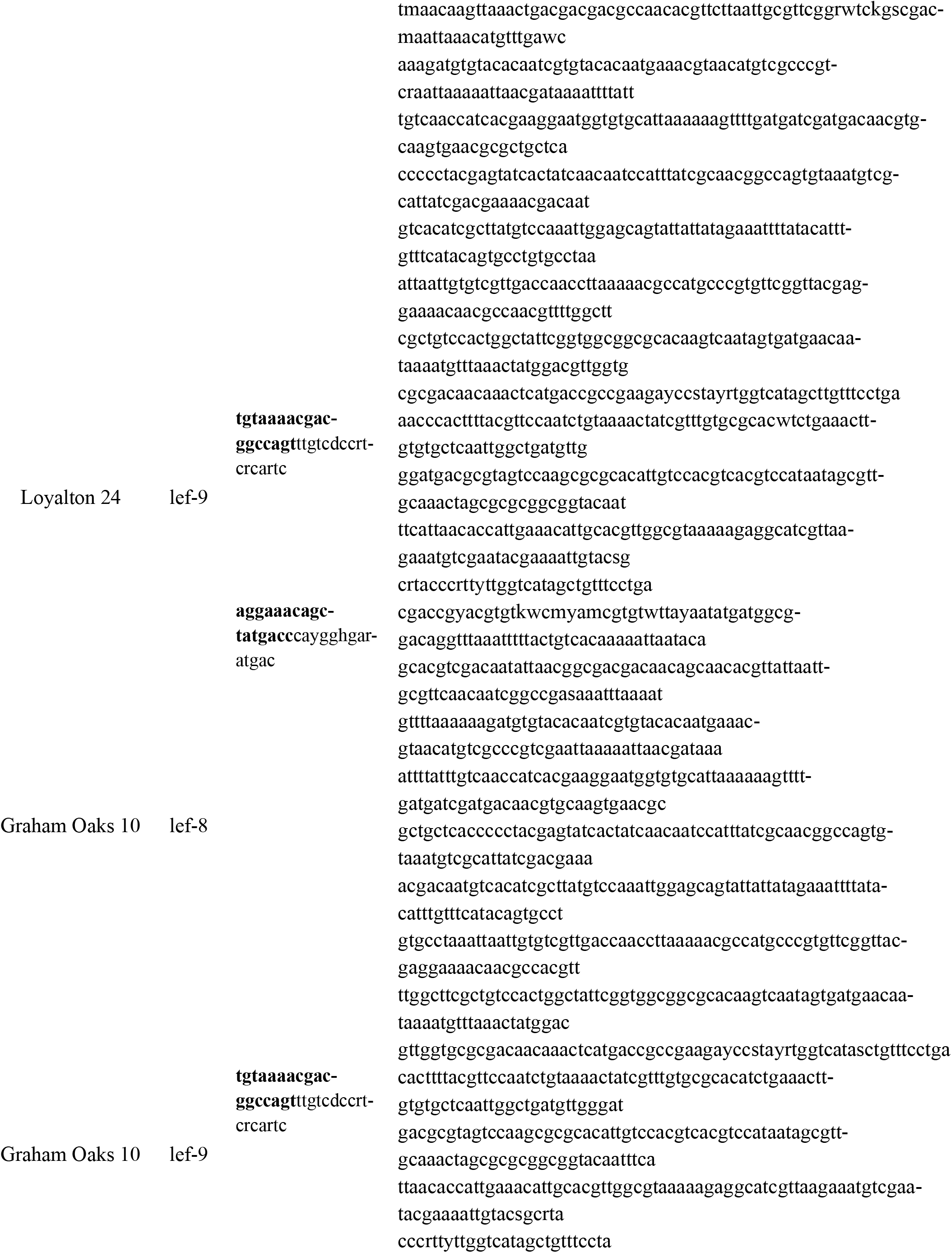
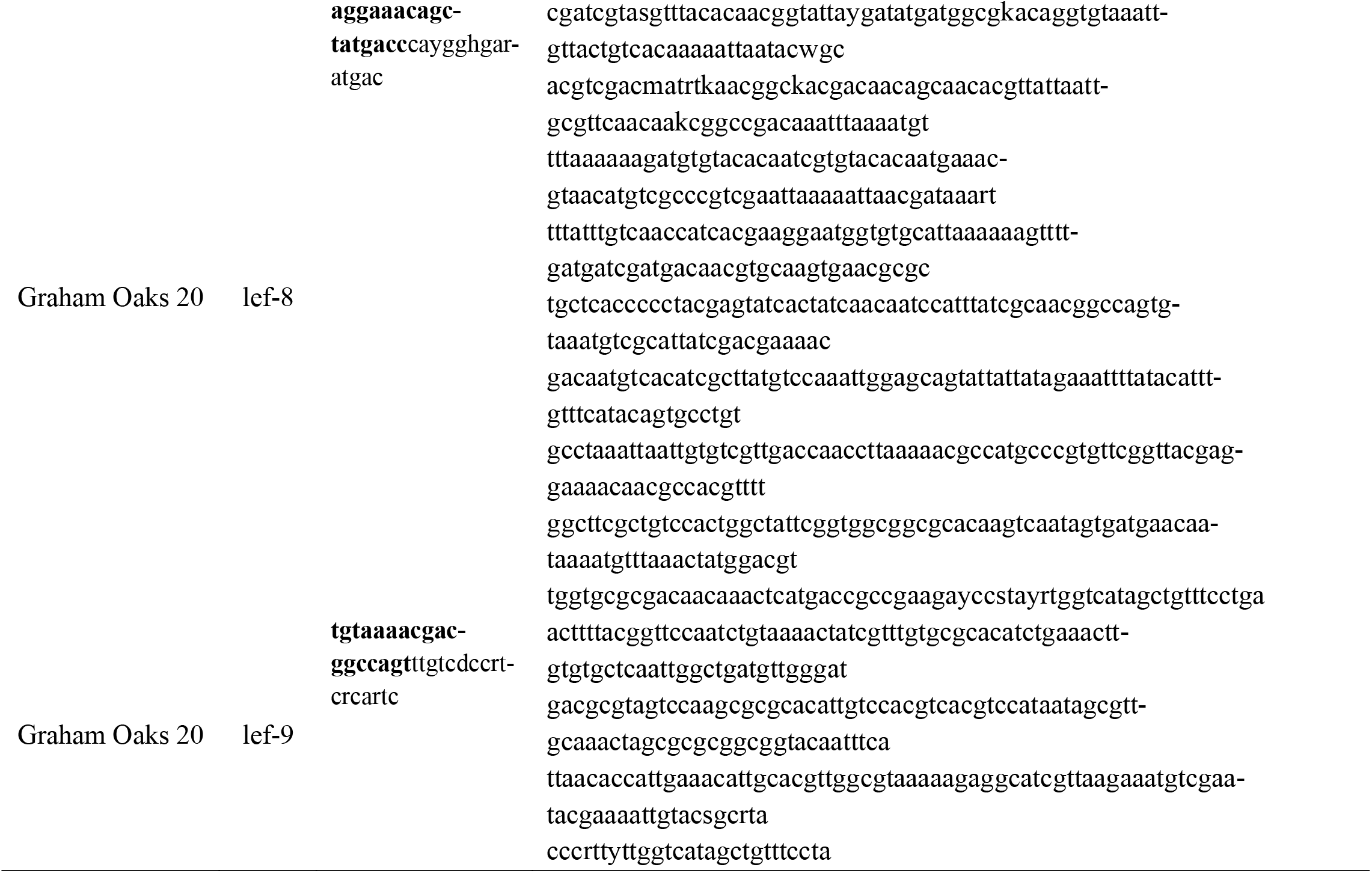
Primers and sequences for viral genes amplified from infected caterpillars

**Figure S1.**
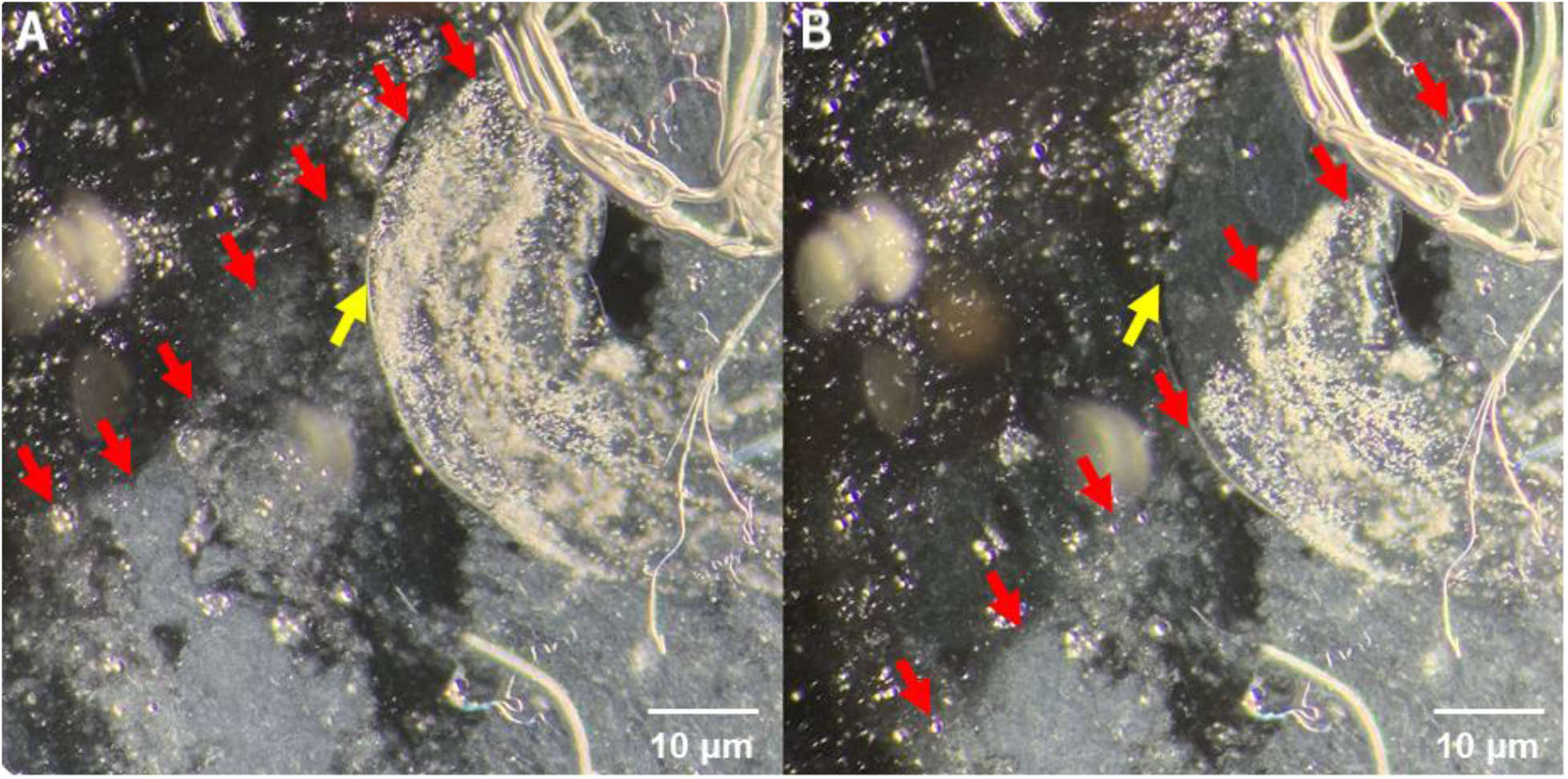
Fat body tissue smear from a caterpillar infected with GV under a light microscope at 200x magnification with phase contrast. A drop of NaOH was added which moved through the sample. The red arrows point to the boundary between NaOH solution and water. The yellow arrows point to the OBs (Figure A) which turned transparent (Figure B) after they were dissolved by the strong base.

**Figure S2.**
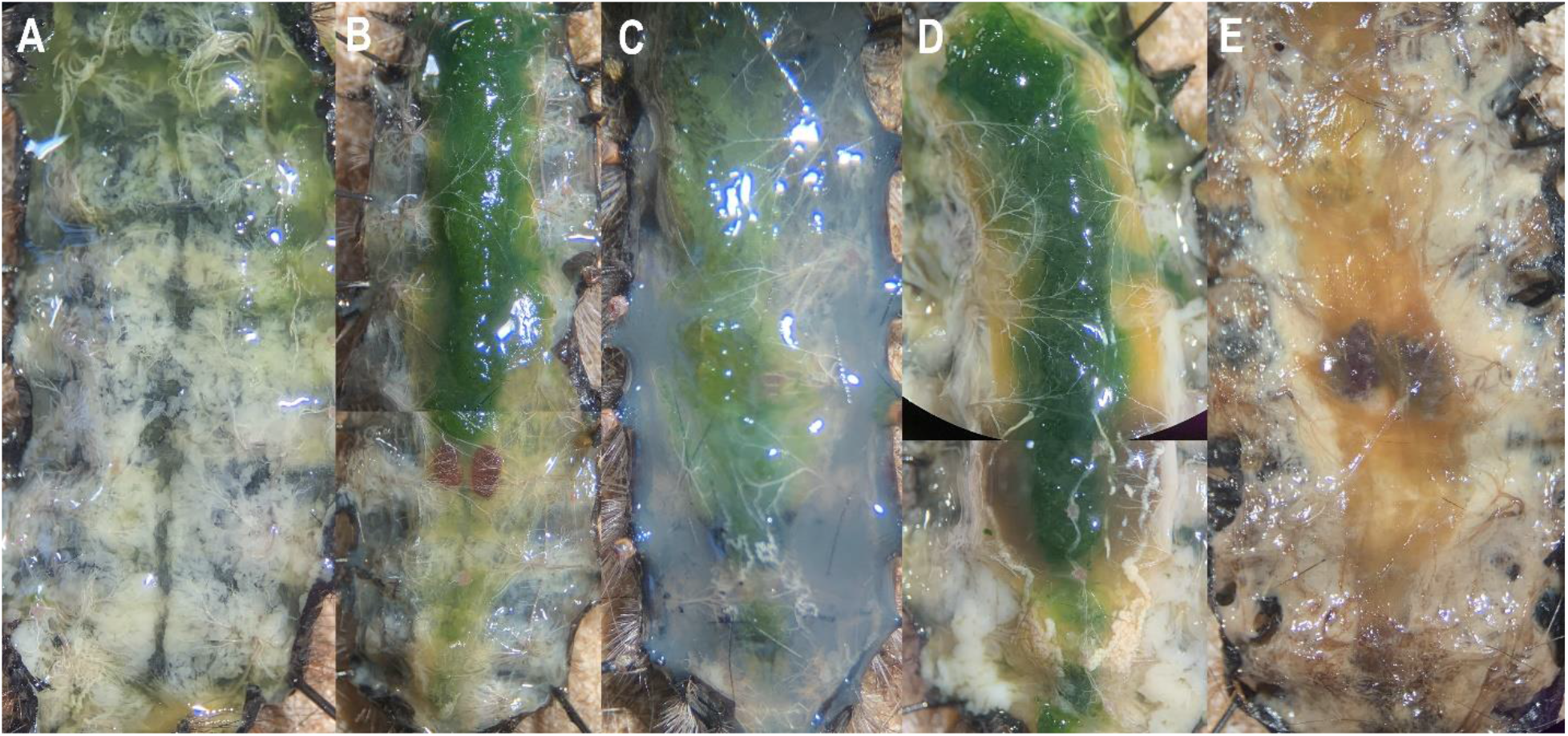
Ventral dissection of late instar *A. virginalis* larva in the order of infection severity, ranging from (A) uninfected, (B) low severity, (C) medium severity, (D) high severity, to (E) very high severity. The caterpillar’s gut in figure A is missing. Caterpillars with higher infection severity have a cloudier hemolymph and a darker and more yellowish fat body. They also often have large clusters of infected tissues next to the midgut (e.g. Figure D).

